# Negative selection on human genes causing severe inborn errors depends on disease outcome and both the mode and mechanism of inheritance

**DOI:** 10.1101/2020.02.07.938894

**Authors:** Franck Rapaport, Bertrand Boisson, Anne Gregor, Vivien Béziat, Stephanie Boisson-Dupuis, Jacinta Bustamante, Emmanuelle Jouanguy, Anne Puel, Jeremie Rosain, Qian Zhang, Shen-Ying Zhang, Joseph G. Gleeson, Lluis Quintana-Murci, Jean-Laurent Casanova, Laurent Abel, Etienne Patin

## Abstract

**Background:** Genetic variants underlying severe diseases are less likely to be transmitted to the next generation, and are thus gradually and selectively eliminated from the population through negative selection. Here, we study the determinants of this evolutionary process in genes underlying severe diseases in humans.

**Results:** We propose a novel approach, CoNeS, integrating known negative selection scores through principal component projection. We compare evidence for negative selection at 319 genes underlying inborn errors of immunity (IEI), which are life-threatening monogenic disorders. We find that genes underlying autosomal dominant (AD) or X-linked IEI are under stronger negative selection than those underlying autosomal recessive (AR) IEI, which are under no stronger selection than genes not known to be disease-causing. However, we find that genes with mutations causing AR IEI that are lethal before reproductive maturity and that display complete penetrance are under stronger negative selection than other genes underlying AR IEI. We also find that genes underlying AD IEI by haploinsufficiency are under stronger negative selection than other genes underlying AD IEI. Finally, we replicate these results in a study of 1,140 genes causing inborn errors of neurodevelopment.

**Conclusions:** These findings collectively show that the clinical outcomes of inborn errors, together with the mode and mechanism of inheritance of these errors, determine the strength of negative selection acting on severe disease-causing genes. These findings suggest that estimating the intensity of negative selection with CoNeS may facilitate the selection of candidate genes in patients suspected to carry an inborn error.

## Background

Negative (or purifying) selection is the natural process by which deleterious alleles are selectively purged from the population (1). In diploid species, the strength of negative selection at a given locus is predicted to increase with decreasing fitness and increasing dominance of the traits controlled by the locus concerned: genetic variants causing early death in the heterozygous state are the least likely to be transmitted to the next generation, as their carriers have fewer offspring than non-carriers (2). Variants of human genes that cause severe diseases are, thus, expected to be the primary targets of negative selection, particularly for diseases affecting heterozygous individuals. Human genes have been ranked according to their levels of negative selection (3–5). Nevertheless, the extent to which negative selection affects known human disease-causing genes, and the factors determining its strength, remain largely unknown, particularly because our knowledge of the severity, mode and mechanism of inheritance of human diseases remains incomplete (3, 6–8). The determinants of negative selection remain largely unknown.

The strength of negative selection at a given gene is classically estimated by comparing the coding sequence of the gene in a given species with that of one or several closely related species; it depends on the proportion of amino-acid changes that have accumulated during evolution (9–11). With the advent of high-throughput sequencing, novel intraspecies estimators have been developed, based on the comparison of the predicted probability of loss-of-function (LOF) mutation for a gene under a random model with the frequency of LOF mutations observed in population databases (5, 12, 13), which capture the species-specific evolution of genes. Using an interspecies-based method and a hand-curated version of the Online Mendelian Inheritance in Man (hOMIM) database, a previous study elegantly showed that most human genes for which mutations cause highly penetrant diseases, including autosomal dominant (AD) diseases in particular, are under stronger purifying selection than genes associated with complex disorders (6). However, other studies based on OMIM genes have reported conflicting results (3, 14–17), probably due to the incompleteness and heterogeneity of the database. Moreover, no study has yet addressed this problem with intraspecies estimators, even though it has been suggested that the choice of the reference species for interspecies estimators contributes to discrepancies across studies (6).

We aimed to improve the identification of determinants of negative selection acting on human disease-causing genes, by developing a new negative selection score combining several informative intra- and interspecies statistics, in a study focusing on inborn errors of immunity (IEI). IEI, previously known as primary immunodeficiencies (PID) (18), are genetic diseases that disrupt the development or function of human immunity. They form a large group of genetic diseases that has been widely studied and they are well-characterized physiologically (immunologically) and phenotypically (clinically) (19, 20). IEI are often symptomatic in early childhood and, at least until the turn of the 20^th^ century and the introduction of antibiotics, most individuals with IEI probably failed to reach reproductive maturity. This suggests that IEI genes were under negative selection from the dawn of mankind until very recently. In this study, we investigated whether the severity of IEI and their mode and mechanism of inheritance have left signatures of negative selection of various intensities in the human genome. We also tested our model on genes underlying inborn errors of neurodevelopment (IEND), another group of well-characterized severe genetic diseases.

## Results

### CoNeS is a novel consensus-based method for measuring negative selection

We developed a new measurement, consensus negative selection (CoNeS), to take into account information from both interspecies (the *f* parameter from SnIPRE (11), lofTool (13) and evoTol (21)) and intraspecies (RVIS (5), LOEUF (22), pLI (12) and SIS (23)) statistics measuring the strength of negative selection. The correlation between these different statistics is shown in figure S1. We excluded from the computation gene-level metrics that do not explicitly measure the strength of negative selection (such as the GDI (24) or pRecessive (12)) or that were unavailable for more than 25% of the genes (such as Sel (3)). CoNeS was obtained through a standardized (i.e. mean of 0 and SD of 1) projection of these seven methods on the first principal component, which captures 81.8% of the total variance. The CoNeS distribution for 17,918 autosomal genes is shown in figure 1a. The distribution is bimodal due to the inclusion of bimodal measurements (pLI and LOEUF) in the calculation. Low CoNeS values are associated with strong selection constraints (i.e. low *f*, lofTool, evoTol, LOEUF and RVIS; high SIS and pLI). As expected, CoNeS values were significantly lower for the X chromosome than for the autosomes (median −0.725, Wilcoxon one-tailed test *P* = 3.04×10^-52^; figure S2), as negative selection acts on deleterious recessive variants in both homozygous females and hemizygous males. We therefore considered X-chromosome genes separately from autosomal genes in all subsequent analyses. We assessed the sensitivity to each of the individual scores, by calculating CoNeS after the removal of each of the seven measurements contributing to the combined score. The resulting scores were strongly correlated with CoNeS (0.947 < Spearman’s *R*^2^ < 0.993; figure S3), indicating that CoNeS is not dependent on a single statistic.

**Figure 1:**
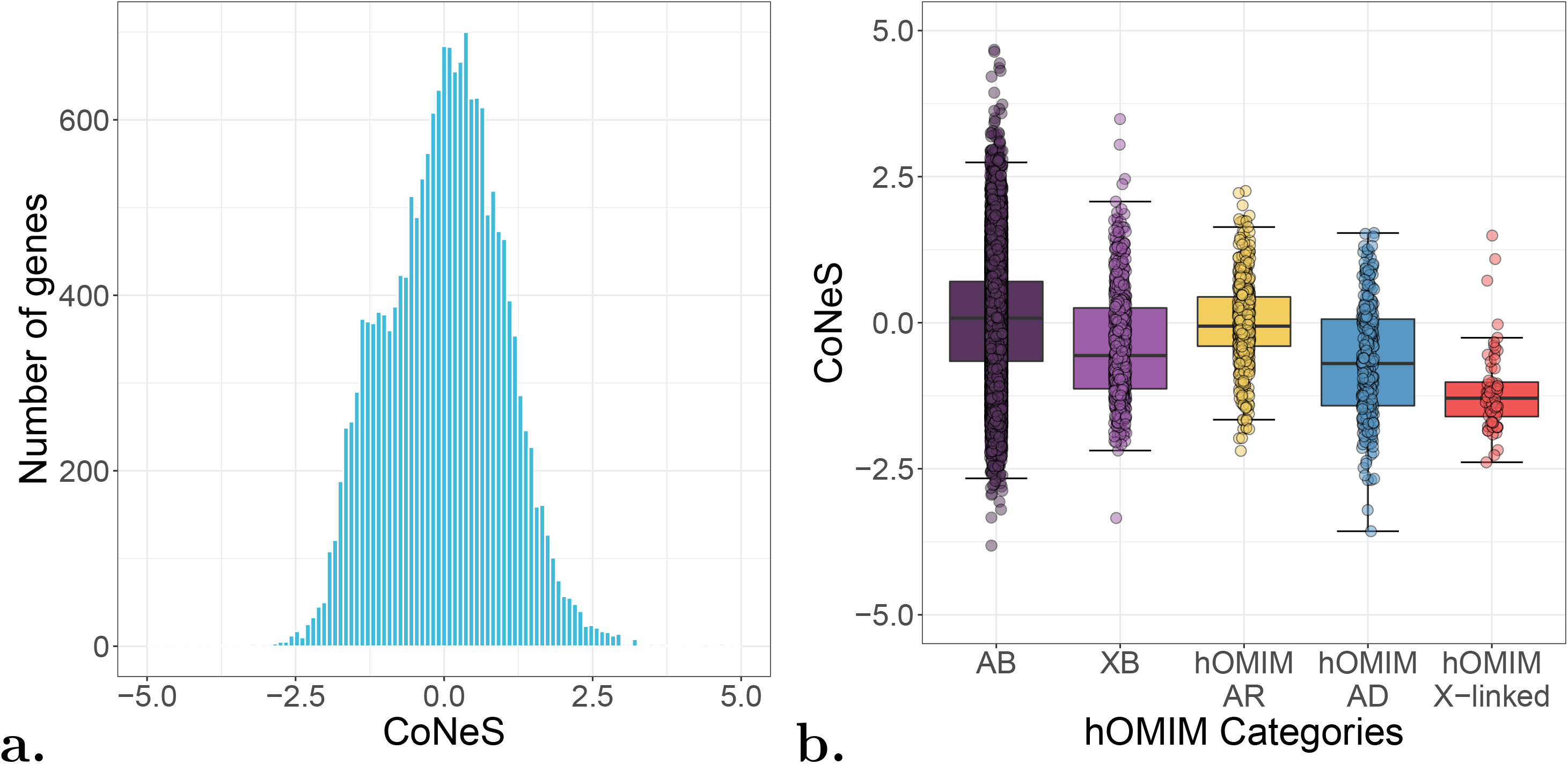
The distribution of CoNeS across human genes. **a.** The distribution of CoNeS across 17,918 autosomal human genes. **b.** The distribution of CoNeS across genes causing Mendelian diseases with complete penetrance (hOMIM), according to their dominant (AD), recessive (AR), or X-linked mode of inheritance, relative to autosomal (AB) and X-chromosome (XB) background genes.

### CoNeS is lower for Mendelian disease-causing genes than for background genes

For the validation of our approach, we sought to replicate previous observations based on the hOMIM database (6). We compared the CoNeS of hOMIM Mendelian disease-causing autosomal and X-chromosome genes to that of 15,166 “autosomal background” (AB) and 604 “X background” (XB) genes, respectively, these background genes being not known to be essential nor to be involved in any severe genetic disorder (see Methods for details). The CoNeS was significantly lower for hOMIM autosomal genes than for the AB group (Wilcoxon one-tailed test: *P* = 1.18×10^-15^; resampling test: *P* < 10^-5^; table 1, fig. 1b), indicating that the hOMIM genes were subject to stronger selection constraints. We replicated a strong effect of the mode of disease inheritance on the strength of negative selection: the difference in CoNeS between hOMIM genes causing AD diseases and the AB group was highly significant (Wilcoxon one-tailed test: *P* = 2.05×10^-31^; resampling test: *P* < 10^-5^; table S1), whereas this difference was not significant for hOMIM genes causing autosomal recessive (AR) diseases (table S2) (fig. 1b). Furthermore, X-linked hOMIM genes were under significantly stronger negative selection (median: −1.29) than XB genes (Wilcoxon one-tailed test: *P* = 1.08×10^-12^; resampling test: *P* < 10^-5^; table S3, fig. 1b). Interestingly, CoNeS separated AD and X-linked hOMIM genes from background genes more effectively than any of the individual scores, with the exception of lofTool and evoTol. Overall, these results show that CoNeS is a valid measurement of the strength of negative selection. They also show that genes underlying known monogenic disorders, especially for AD and X-linked disorders, are under stronger selection constraints than the rest of the coding genome.

**Table 1:** Statistical significance of differences in negative selection scores between genes causing autosomal dominant (AD) inborn errors of immunity (IEI) and autosomal background (AB) genes. The P-values for a one-tailed Wilcoxon test and a resampling-based test (Methods) assessing the difference between IEI AD genes and AB genes are shown.

| Score | Wilcoxon-based test | Resampling-based test |
| --- | --- | --- |
| evoTol | 0.425 | 0.767 |
| *f* parameter | $1.32 \times 10^{-7}$ | $5 \times 10^{-4}$ |
| LOEUF | $1.75 \times 10^{-7}$ | $< 10^{-5}$ |
| lofTool | $1.08 \times 10^{-4}$ | $2.70 \times 10^{-3}$ |
| pLI | $8.99 \times 10^{-6}$ | $< 10^{-5}$ |
| RVIS | $2.21 \times 10^{-5}$ | $2.42 \times 10^{-2}$ |
| SIS | $4.36 \times 10^{-5}$ | $7.60 \times 10^{-3}$ |
| CoNeS | $7.58 \times 10^{-9}$ | $< 10^{-5}$ |

### The mode of inheritance strongly affects the strength of negative selection on IEI genes

We then focused on autosomal IEI genes. There are 359 known IEI, caused by defects of 319 genes, in the latest IUIS committee classification (18). Historically, IEI were considered to be Mendelian disorders, with both complete clinical penetrance and detectable immunological abnormalities. More recently, IEI with incomplete penetrance and/or without detectable immunological phenotypes have been described (19). More than two thirds of the known IEI genes (224/319) cause IEI that are AR; a smaller number of genes (51/319) cause IEI that are AD; an even smaller number cause IEI that are X-recessive (XR) (19/319), and one unique gene (WAS) causes an IEI that is X-dominant (XD). A small number of IEI-causing loci cause diseases with both AR and AD inheritance patterns (24/319) (fig. 2a). Consistent with the results obtained for hOMIM genes, the CoNeS for IEI AD genes was significantly lower than that for random groups of genes of similar length (median: −1.14, Wilcoxon test *P* = 7.58×10^-9^ resampling test *P* < 10^-5^), whereas that for IEI AR and IEI AD/AR genes was not (medians: –0.045 and −0.211, resampling-test *P* = 0.180 and *P* = 0.172 respectively) (fig. 2b, tables 1, S4 and S5). Most individual statistics (with the exception of evoTol) showed IEI AD genes to be under significantly stronger negative selection than the genes of the AB group, but the difference was the most significant for CoNeS (table 1). These results demonstrate that genes causing AD IEI are under stronger negative selection than genes causing AR IEI or genes causing both AR and AD IEI, consistent with the notion that the dominance of the disease has a strong impact on the strength of negative selection on human disease-causing genes.

**Figure 2:**
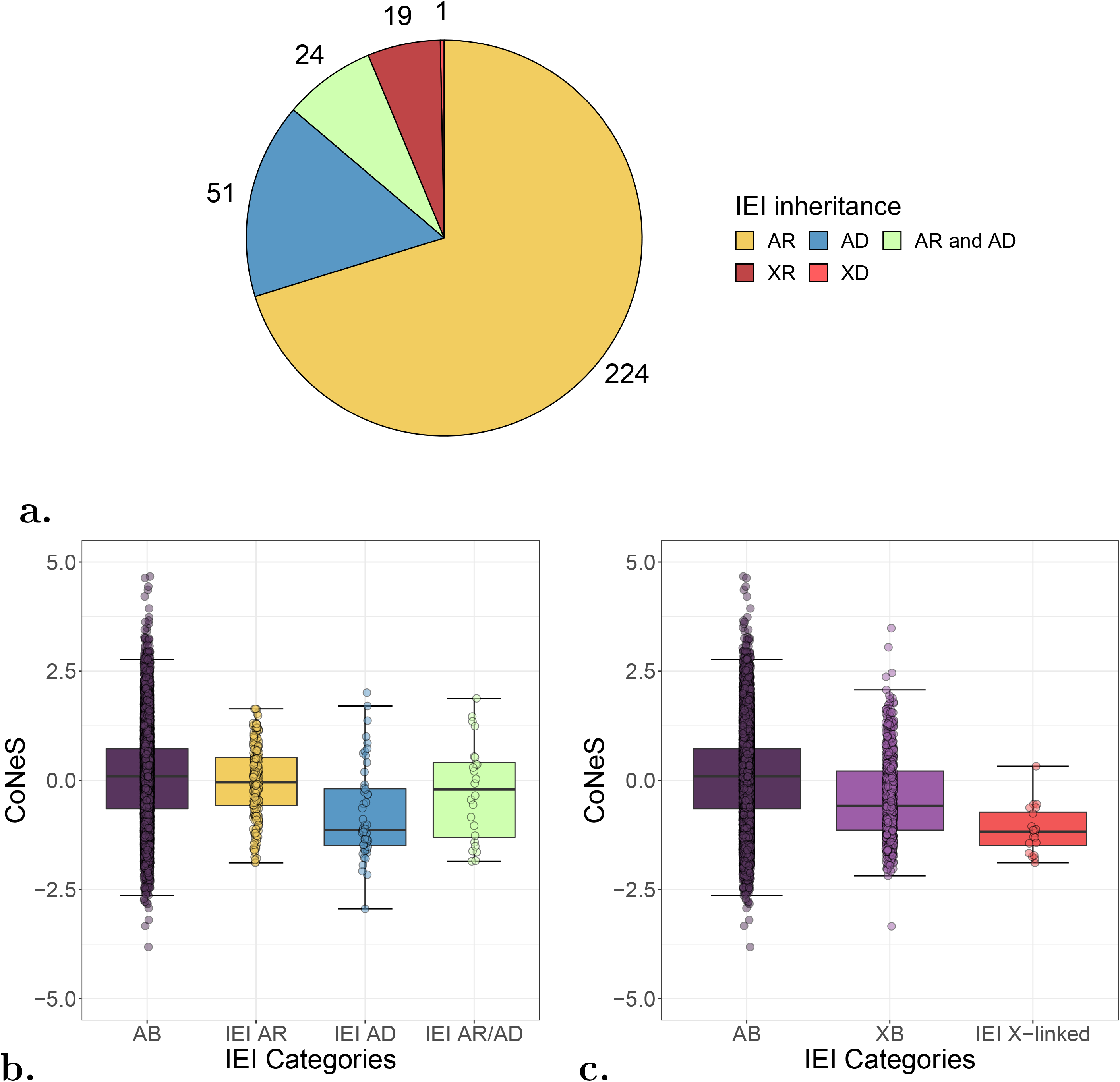
The distribution of CoNeS across genes causing inborn errors of immunity (IEI). **a.** The number of genes for each of the mode of inheritance of the IEI genes. **b.** The distribution of CoNeS across autosomal genes causing IEI, according to their dominant (AD), recessive (AR), or both dominant and recessive (AR/AD) mode of inheritance, relative to autosomal background (AB) genes. **c.** The distribution of CoNeS on the X-linked genes causing IEI, relative to autosomal (AB) and X-chromosome (XB) background genes.

### X-linked IEI-causing genes are under stronger negative selection than other X-linked genes

Males carry only one copy of the X chromosome. The mutations underlying XR diseases would therefore be expected to be purged from the population more rapidly, and to be under stronger selection constraints than the mutations underlying AR diseases. Consistent with this hypothesis, we found that the CoNeS for XB gene was significantly lower than that for AB genes (fig 2c). IEI XR genes were under even stronger selection constraints than XB genes, as indicated by their lower CoNeS (median −1.17, Wilcoxon test: *P* = 1.19×10^-5^, resampling-based test: *P* = 2.27×10^-2^, table S6). CoNeS was one of the statistics yielding the most significant difference between IEI XR and XB, together with pLI (table S6). At individual gene level, only one IEI XR gene had a positive CoNeS value: *CSF2RA* (0.324). Human *CSF2RA* deficiency causes juvenile pulmonary alveolar proteinosis, a disease that was lethal until very recently (25). However, *CSF2RA* lies in the pseudo-autosomal region of the X and Y chromosomes, so heterozygous males do not develop the disease. Serving as a natural control, this gene was, therefore, unsurprisingly under weaker selection than the other XR IEI genes. The CoNeS results show that IEI-causing genes on the X chromosome are, like autosomal IEI-causing genes, under stronger selective constraint than the rest of the coding genome.

### Disease severity increases the strength of negative selection acting on IEI AR genes

We investigated whether the disease genes that decrease fitness the most were under the strongest selective constraints, by classifying IEI-causing genes into two categories: 213 “high-severity” genes that, when mutated, cause severe disease and prevent patients from reaching reproductive age in the absence of modern treatment, and 104 “mild-severity” genes, comprising all the other genes causing diseases with incomplete penetrance and/or a more moderate impact, as demonstrated by the findings for at least one reported multigenerational multiplex family (whether dominant or recessive). AR and XR IEI disease-causing genes are enriched in “high-severity” genes (80.1% and 75.0%, respectively), whereas AD diseases are typically associated with “mild-severity” genes (77.6%) (χ^2^ test *P* = 8.06×10^-15^; fig. 3a). This observation suggests that recessive mutations decrease fitness more than dominant mutations, consistent with the negative relationship observed between fitness and the dominance coefficient in *Drosophila*, yeast, and thale cress (26–28). However, caution is required, because this enrichment may also be due to a bias in the IEI database (e.g., dominant, severe diseases are more difficult to study). Interestingly, we observed that the IEI AR genes of the “high-severity” group were under significantly stronger negative selection than those of the “mild-severity” group (medians – 0.0833 and 0.495 respectively, Wilcoxon test: *P* = 8.27×10^-5^) or AB genes (Wilcoxon one-tailed test: *P* = 1.65×10^-3^, table S7). Disease severity did not significantly influence CoNeS for IEI AD, IEI AR/AD and IEI XR genes (Wilcoxon one-tail test *P* = 0.415, 0.408 and 0.433 respectively see fig. 3b), although this finding may reflect lack of power. Together, our findings show that genes causing severe AR IEI are under stronger selective constraints than genes causing milder AR IEI, providing direct evidence that disease severity affects the strength of negative selection acting on human disease-causing genes.

**Figure 3:**
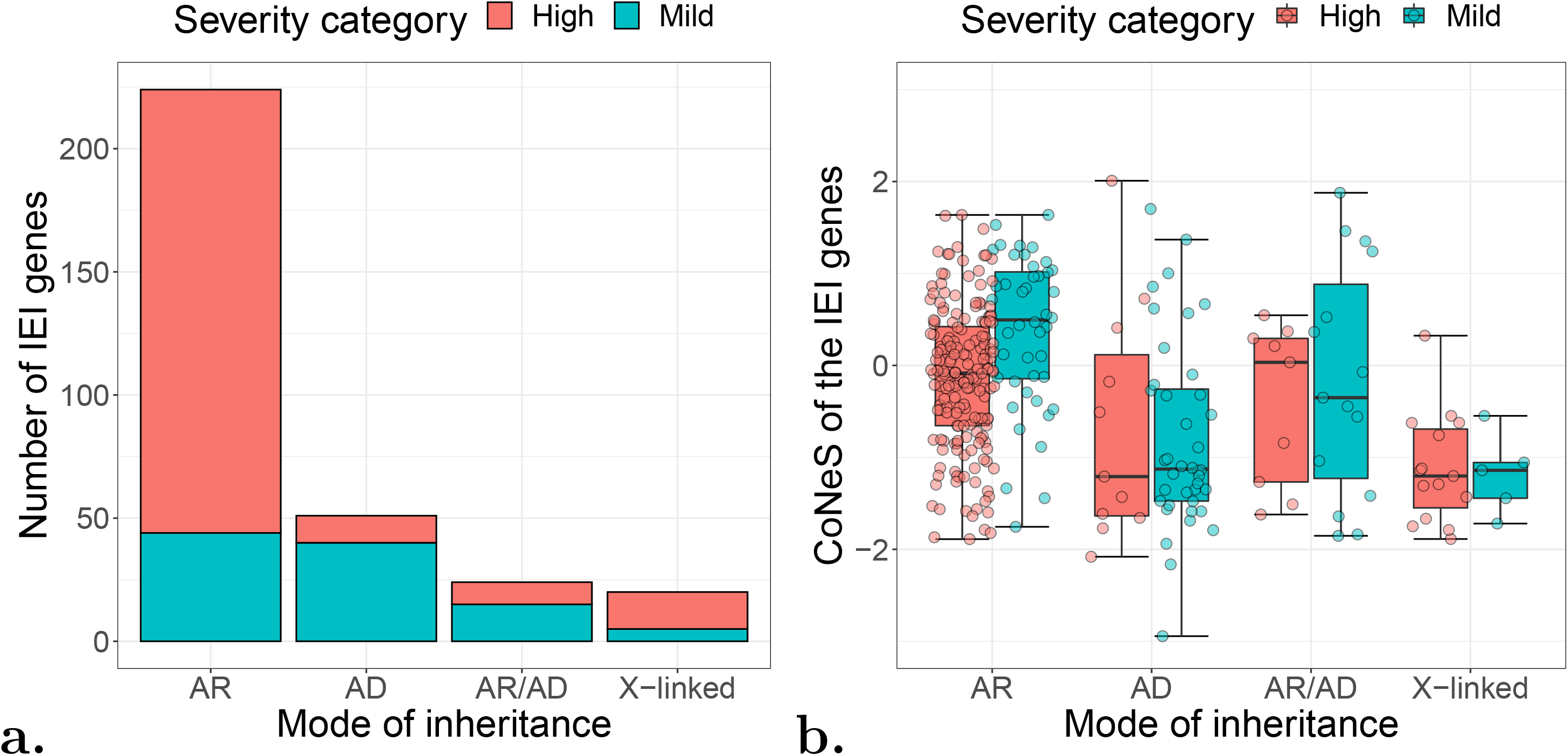
The distribution of CoNeS for genes causing inborn errors of immunity (IEI), according to disease mode of inheritance and clinical severity. **a.** The number of IEI genes for each severity category and each mode of inheritance. **b.** The distribution of CoNeS for the IEI genes, as a function of phenotype severity and mode of inheritance.

### Haploinsufficient genes are under stronger selection than dominant negative or gain-of-function genes

Dominance can operate by negative dominance, haploinsufficiency (HI), or gain-of-function (GOF) (29). In AD disorders due to dominant negative (DN) alleles, the AD cellular and clinical deficiencies are caused by the interference of the mutant gene product with the activity of the wild-type (WT) product, whatever the molecular mechanism. In AD disorders due to HI, the mutant copy is not functional and does not interfere with the WT product, and the single functional WT copy produces too little protein to fulfill its function. HI is more commonly associated with loss-of-expression alleles and DN with normally or highly expressed alleles, but rare examples have been reported of HI with normal levels of the mutant protein (30), and of negative dominance (ND) with a lack of detectable mutant protein (31). Autosomal dominance by GOF defines a third category, in which the mutant protein is produced (32). We hypothesized that genes causing disease through HI mechanisms are under stronger negative selection than those in which the underlying mechanism is ND or GOF, because any loss-of-expression mutation at HI loci is likely to be LOF and potentially disease-causing (33). Dominant forms of IEI have been reported to be caused by variants acting by HI (20 genes), GOF (14 genes), and ND (9 genes) (supplementary files). *RAC2* is the only AD gene to have been shown to be associated with two different mechanisms (ND and GOF) and *TLR3* with two similar mechanisms (HI and ND) (34); we classified both these genes as having “unknown” modes of dominance. As expected, the CoNeS values of genes operating by HI were lower than those of DN and GOF genes (medians of −1.35, −0.300 and −0.357, respectively) (fig. 4). Despite the small number of genes in each group, the difference between the HI group and the group containing both DN and GOF genes was statistically significant (Wilcoxon one-tailed test: *P* = 7.34×10^-4^, table 2). The only individual statistics that separated these groups were pLI and LOEUF, which explicitly measure the strength of selection on heterozygotes (22, 35) and are therefore better suited to this specific task than other scores (table S8). Caution is also required because statistics based on LOF mutations (LOEUF, pLI, RVIS and LoFTool) do not depart from genome-wide expectations for GOF genes, probably because these statistics typically ignore GOF alleles when scoring genes. Thus, the mechanism of dominance (HI, DN, or GOF) affects the strength of negative selection acting on human genes causing AD IEI.

**Figure 4:**
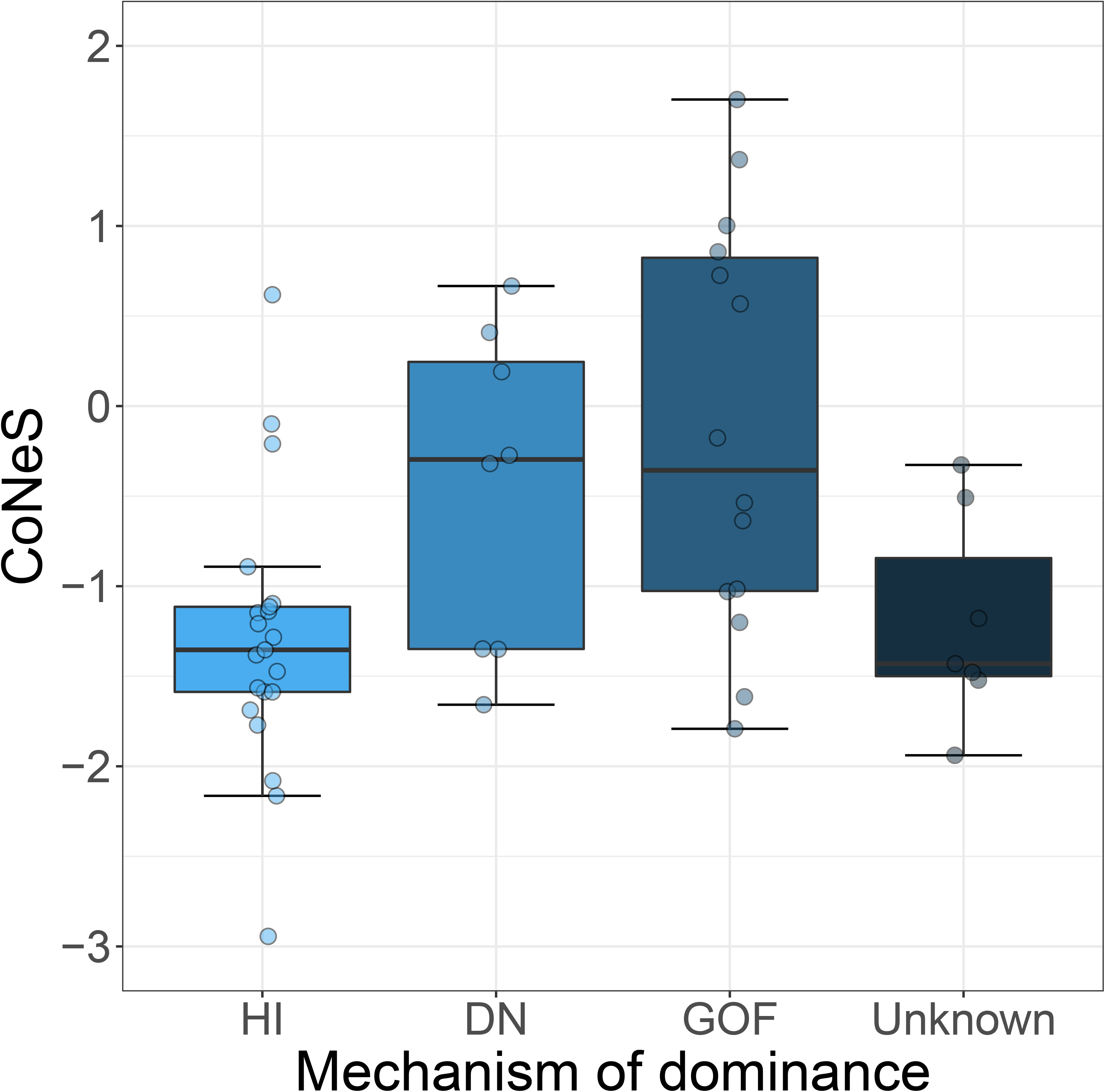
The distribution of CoNeS for the genes causing autosomal dominant inborn errors of immunity according to mechanism of dominance.

**Table 2:** Statistical significance of differences in negative selection scores between genes causing inborn errors of immunity (IEI), according to disease mode of inheritance, clinical severity, gene length and GC content. This table presents, for each of the individual scores, the P-values from a multiple re-gression model that predicts each negative selection individual score for each autosomal IEI gene. Predictors include AD and ARAD, two binary variables that code if the gene causes autosomal dominant (AD) disease or both autosomal recessive (AR) and AD diseases, respectively, disease severity (high or mild), the coding sequence (CDS) length (in bp) and the CDS content in C and G bases (GC-content). Total is the total performance of the linear model.

| Score | AD | ARAD | Severity | CDS length | GC content | Total |
| --- | --- | --- | --- | --- | --- | --- |
| evoTol | 0.71 | 0.43 | $4.8 \times 10^{-2}$ | 0.97 | $3.5 \times 10^{-2}$ | $8.8 \times 10^{-2}$ |
| *f* parameter | $3.4 \times 10^{-9}$ | $4.8 \times 10^{-2}$ | $3.3 \times 10^{-3}$ | 0.68 | 0.11 | $3.9 \times 10^{-7}$ |
| LOEUF | $2.9 \times 10^{-6}$ | 0.18 | $4.7 \times 10^{-3}$ | 0.12 | 0.94 | $1.3 \times 10^{-4}$ |
| lofTool | $5.1 \times 10^{-3}$ | $9.2 \times 10^{-2}$ | $3.3 \times 10^{-2}$ | 0.80 | $3.0 \times 10^{-3}$ | $1.7 \times 10^{-3}$ |
| pLI | $3.9 \times 10^{-9}$ | $5.1 \times 10^{-3}$ | $4.9 \times 10^{-2}$ | 0.29 | 0.72 | $1.3 \times 10^{-7}$ |
| RVIS | $8.6 \times 10^{-4}$ | 0.61 | $9.7 \times 10^{-2}$ | 0.77 | $4.5 \times 10^{-4}$ | $4.0 \times 10^{-4}$ |
| SIS | $6.6 \times 10^{-5}$ | 0.90 | $3.1 \times 10^{-2}$ | 0.62 | 0.41 | $2.3 \times 10^{-3}$ |
| CoNeS | $3.6 \times 10^{-9}$ | $5.8 \times 10^{-2}$ | $2.2 \times 10^{-3}$ | 0.63 | 0.45 | $8.3 \times 10^{-7}$ |

### Disease severity, mode of inheritance, and mechanism of dominance independently affect the strength of negative selection on IEI genes

Recessive IEI tend to be more severe than dominant IEI. We therefore investigated whether dominance and disease severity affected the measured strength of the negative selection acting on IEI-causing genes in an independent manner. We fitted a linear regression model to all IEI autosomal genes, predicting CoNeS and using as covariates the mode of inheritance, the severity of the associated disease, coding sequence length and gene GC content (coded as a percentage) (see tables 3 and S9). This multiple linear regression model predicted CoNeS with significant accuracy (*P* = 8.3×10^-7^). The mode of inheritance and severity predicted CoNeS better than they would predict any of the individual scores, with the exception of pLI, whereas coding sequence length and GC-content did not improve predictive performance (*P* = 0.63 and 0.45, respectively). GC-content significantly improved the prediction of evoTol, lofTool and RVIS (*P* = 3.5×10^-2^, 3.0×10^-3^ and 4.5×10^-4^, respectively), these relationships likely being due to computational artifacts in these methods (see table 3 for a full comparison). Coding sequence length did not improve the prediction score for any method. When IEI AD genes were considered separately, the mechanism of dominance improved prediction even further (ANOVA *P* = 1.76×10^-3^). These results demonstrate that mode of inheritance, mechanism of dominance, and clinical severity of IEI are three independent determinants of the strength of negative selection, as measured by CoNeS.

### The mode of inheritance and mechanism of dominance affect the strength of negative selection on genes causing inborn errors of neurodevelopment

For further validation of the results obtained for IEI genes, we compared the CoNeS values of genes underlying IEND, a group of severe, early-onset diseases with well-characterized genetic etiologies (36). We classified the 1,140 IEND-causing genes according to their mode of inheritance: 650 genes cause AR IEND, 303 cause AD IEND and 65 cause both AR and AD IEND, whereas 46 and 6 X-linked genes cause XR and XD IEND, respectively, and 70 X-linked IEND genes have an unknown mode of inheritance (fig. 5a). Consistent with the results obtained for IEI, AD IEND genes were found to be under stronger negative selection (median −1.61) than AB genes (Wilcoxon one-tailed test: *P* = 4.54×10^-133^ and resampling test: *P* <10^-5^) (fig 5b, table S10) whereas AR IEND genes were found not to be subject to strong selection (median −1.44 ×10^-2^, Wilcoxon one-tailed test: *P* = 3.69×10^-4^, resampling-based *P* = 0.192, table S11). By contrast to the IEI results, AR/AD IEND genes were found to be under stronger negative selection than AB genes (median −1.01, Wilcoxon test *P* = 4.55×10^-14^, table S12 and fig. 5b), although to a lesser degree than IEND AD genes. X-linked IEND genes were also found to be under stronger selective constraints than XB genes (Wilcoxon one-tailed test: *P* = 3.39×10^-37^; fig 5c, table S13). We found no significant difference between the different modes of transmission via the X chromosome (medians −1.48, −1.43 and −1.54 for the XD, XR and X unknown modes of inheritance, respectively). Finally, CoNeS was lower for the 237 IEND AD acting via HI (median −1.69), than for the 44 IEND AD genes not acting via HI (median −0.857) (*P* = 1.12×10^-13^) (fig 5d, table S14). These results confirm that the strength of negative selection acting on genes causing severe diseases depends on the mode of inheritance and mechanism of dominance of the disease.

**Figure 5:**
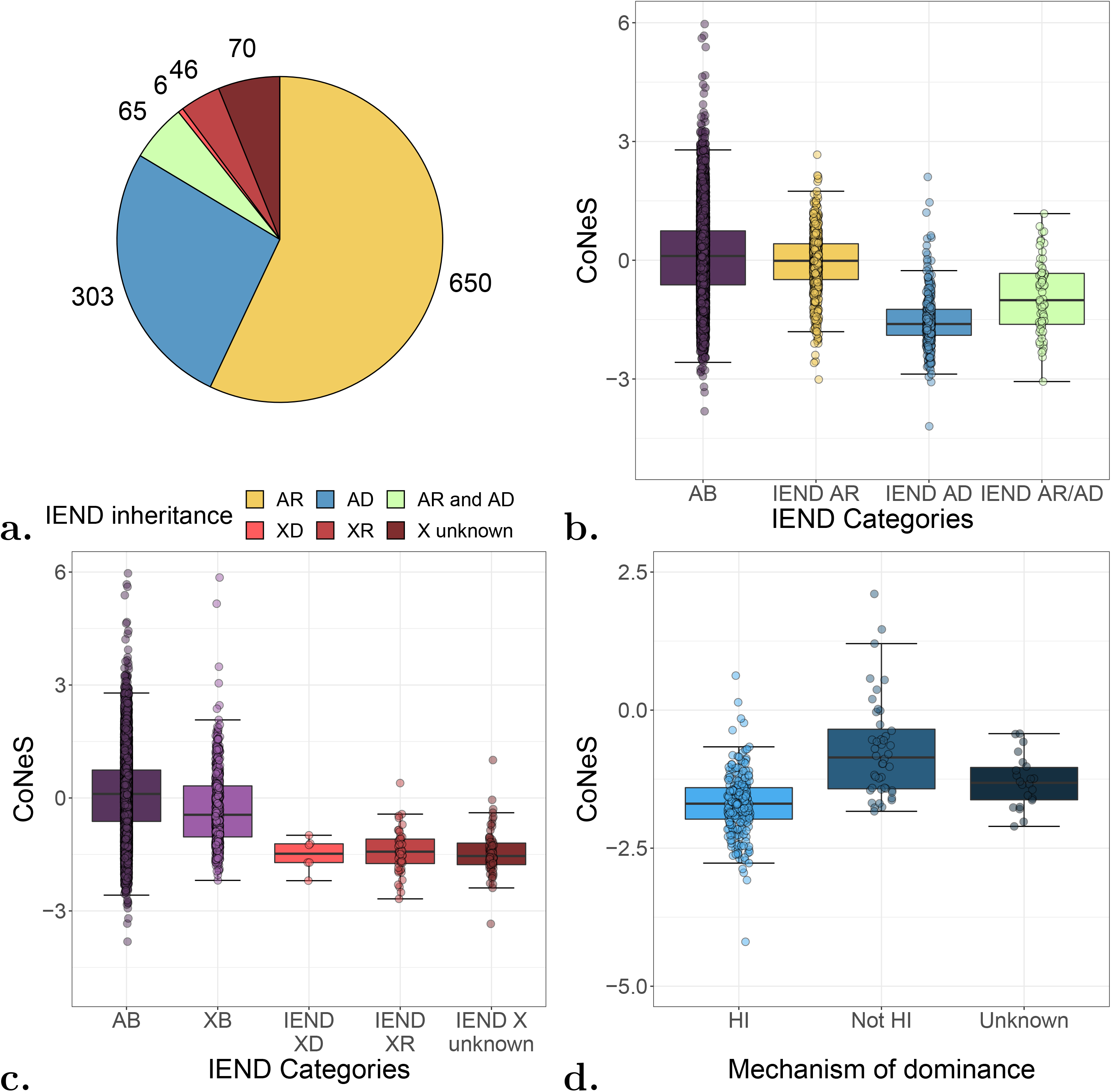
The distribution of CoNeS for genes causing inborn errors of neurodevelopment (IEND), according to disease mode of inheritance and mechanism of dominance. **a.** The number of genes for each of the mode of inheritance of the IEND genes. **b.** The distribution of CoNeS across autosomal genes causing IEND, according to their dominant (AD), recessive (AR), or both recessive and dominant (AR/AD) mode of inheritance, relative to autosomal background (AB) genes. **c.** The distribution of CoNeS on the X chromosome, relative to AB genes. **d.** The distribution of CoNeS for the IEND AD genes according to the mechanism of dominance.

## Discussion

We have developed a novel score that combines inter- and intraspecies measurements of negative selection to determine the mechanisms affecting the strength of this selection at human disease-causing loci. We first focused on genes responsible for IEI, and then validated our results with genes causing IEND. Both these groups of genetic diseases are severe and thoroughly annotated. We demonstrated that the CoNeS score accurately estimated the strength of negative selection acting on human genes, as it was more negative for genes on the X chromosome than for those on autosomes, for loci causing Mendelian diseases than for other genes, and at loci causing dominant diseases than at loci causing recessive diseases. Importantly, we showed that the CoNeS score was significantly more negative at loci causing recessive IEI with a high degree of clinical severity than for other recessive IEI-causing genes. This result contrasts with the nonsignificant difference observed in a previous study based on highly penetrant OMIM genes (6), but confirms a more recent study based on the Bayesian estimation of the selection coefficient of heterozygotes for haploinsufficient genes (8). This discrepancy probably reflects differences in power and in the annotations of disease databases. Together, these results indicate that the effects of negative selection on genetic variation depend on both the mode of inheritance and the clinical outcome of human diseases.

This study provides the first evidence, to our knowledge, that genes causing AD diseases by haploinsufficiency are under stronger negative selection than other AD genes. The significantly more negative CoNeS score for HI genes than for other AD genes can be accounted for principally by the inclusion of the LOEUF and pLI statistics (tables S8 and S14). pLI was originally described as “the probability of being loss-of-function intolerant” and has been used for the explicit classification of HI genes (12). However, it was recently argued that pLI cannot be used to infer the HI status of genes directly, because it reflects only the strength of selection acting on heterozygotes (35). Here, we show that both pLI and CoNeS differ significantly between HI and non-HI genes responsible for severe AD diseases, probably because pLI is a measurement of the strength of negative selection and negative selection acts more strongly on HI genes. For a gene acting by HI, any LOF mutation may affect expression levels, thereby disrupting the molecular function of the protein, whereas most mutations of GOF or DN genes will leave the function of the protein intact (33). We show that this observation can translate quantitatively into stronger evolutionary constraints on genes acting via HI than on other autosomal genes underlying dominant conditions.

One of the limitations of our study is the assumption that all mutations at a given locus cause diseases with the same severity and mode of inheritance, and that negative selection is constant within genes. Several genes, such as *C3* (37) and *STAT1* (38), were found to be under strong negative selection (CoNeS of −1.64 and −1.84, respectively), but associated with several diseases of different severities, modes of inheritance and/or incomplete penetrance. The additive effects of multiple small constraints on most of the sequence result in strong overall constraints on these genes. Conversely, a small number of IEI AD genes are under weak negative selection, whereas their mutations underlie severe diseases. For instance, mutations of *TCF3* (CoNeS of 0.409) underlie an AD deficiency of the E47 transcription factor (39), a very severe disease. However, all reported patients share an identical mutation in the small bHLH domain of the *TCF3* gene, suggesting that there may be heterogeneity in the selective constraints on the gene. We tested this hypothesis with subRVIS (40), a domain-level version of RVIS (5) (figure S4). Our findings confirmed that most domains of the gene were not particularly constrained (subRVIS = 83.8, i.e., 83.8% of the domains of all human proteins are under stronger constraints), but the bHLH domain was under relatively strong negative selection (subRVIS = 18.4). These examples suggest that future studies should take such heterogeneity into account and integrate local measurements of selective constraints (40, 41).

## Conclusions

Investigators searching for genes and variants underlying monogenic inborn errors have developed various gene- and variant-level methods for selecting candidate loci (42, 43). Our approach provides an additional tool that could be applied to candidate genes for severe genetic diseases other than IEI and IEND. We recommend comparing candidate genes for a given condition with genes underlying diseases with the same mode and mechanism of inheritance. For example, a heterozygous nonsense variation in a gene with a CoNeS similar to that of AR genes is unlikely to cause disease by HI. Conversely, a candidate gene for an AD disease that is under strong purifying selection is a good candidate. By integrating negative selection scores with other gene-specific metrics, such as pathway centrality (44) and epigenetic marks (45), future studies based on supervised machine-learning (46, 47) will help to identify strong candidates for genetic disorders, ultimately facilitating the dissection of the genetic etiologies of human diseases.

## Methods

### Gene and disease annotations

The lists of hOMIM and IEI genes and their modes of inheritance were obtained from previous publications (6, 18). Each IEI gene was manually annotated for severity and (when AD) for mode of dominance (supplementary file 1). The IEND gene list was assembled from the SysID reference database (36) (supplementary file 1). The AB and XB gene groups included all human genes not listed in the OMIM, IEI or IEND lists or in the list of essential mouse and human genes defined in a previous study (48).

### Computation of the scores

EvoTol (21), lofTool (13) and SIS (23) statistics were downloaded from the corresponding publications. Values for pLI (12) and LOEUF (22) were obtained from gnomAD v.2.1. For RVIS (49), we downloaded the values calculated with ExAC v2 from the RVIS website. For the *f* parameter from SNIPRE (11), we used the values calculated in a previous study (50). We unified the gene names through the *checkGeneSymbols* function of the HGNChelper package version 0.8.1 (51). For each of these scores, we computed the missing values with the *imputePCA* function of the missMDA package version 1.14 (52). We then used the *PCA* function from FactoMineR version 1.41 (53) and the first component, which we standardized through the *scale* function, as the CoNeS score. In total, we computed the individual statistics and CoNeS for 18,460 genes (supplementary file 1). For the calculation of subRVIS (40) for *TCF3*, we used the subRVIS website with the options domain-level and quantile values. We used R version 3.5.2.

### Comparison with random groups of genes

For comparisons of negative selection statistics for a test group of autosomal (or X-chromosome) genes with the AB background (or XB) group, we created 100,000 groups of randomly sampled genes with a coding sequence length in the same decile of the genome-wide distribution as those of the test group. *P*-values were estimated as the proportion of random groups with a median for negative selection statistics below that of the test group. Based on the number of random samples, the lowest non-zero *P*-value possible is 1/100,000=10^-5^. For a proportion of 0, we therefore noted P<10^-5^.

## Supplemental Data

Supplemental Data includes four figures, fourteen tables and one.xlsx file.

## Declarations

### Ethics approval and consent to participate

Not applicable

### Consent for publication

Not applicable

### Availability of data and materials

All data generated or analyzed during this study is included in supplementary file 1.

### Authors’ contributions

Conceptualization, F.R., J.L.C. and L.A.; Methodology, F.R., J.L.C., L.A., L.Q.M. and E.P.; Investigation, F.R., B.B., A.G., V.B., S.B.D., J.B., E.J., A.P., J.R., Q.Z., S.Y.Z., J.G.G. and E.P.; Writing - Original Draft, F.R. and J.L.C.; Writing - Review & Editing, F.R., B.B., J.L.C., L.A., L.Q.M. and E.P.; Funding Acquisition, F.R., J.L.C. and L.A.; Resources, J.L.C. and L.A.; Supervision, J.L.C., L.A., L.Q.M. and E.P.

## Acknowledgements

We would like to thank members of the Laboratory of Human Genetics of Infectious Diseases for helpful discussions and critical reading.

