## Supplemental Figures for "Negative selection on human genes causing severe inborn errors depends on disease outcome and both the mode and mechanism of inheritance"

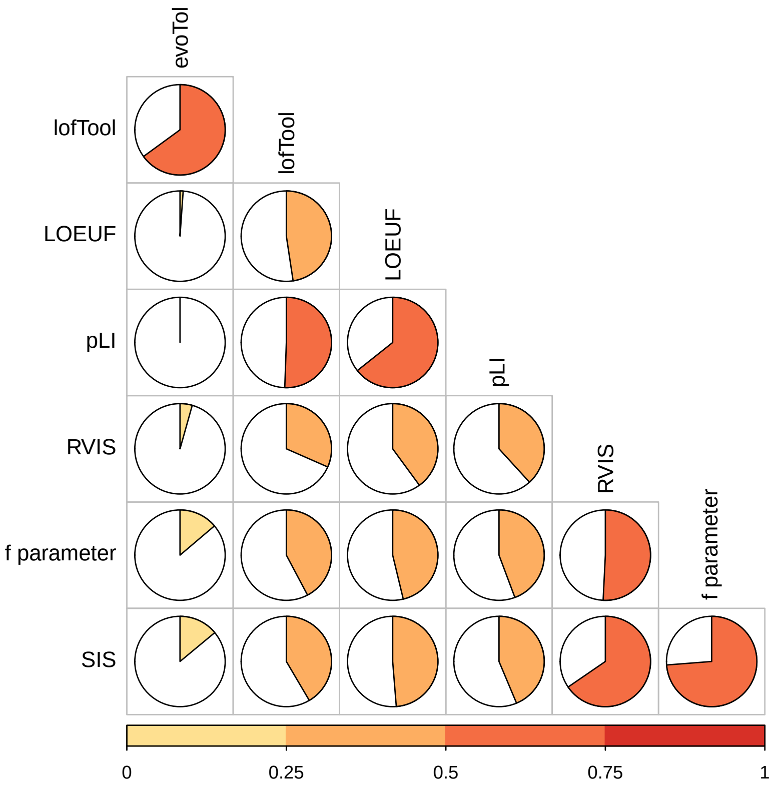


Figure S1: Correlation between the different individual scores, as measured by Spearman's $R^2$.

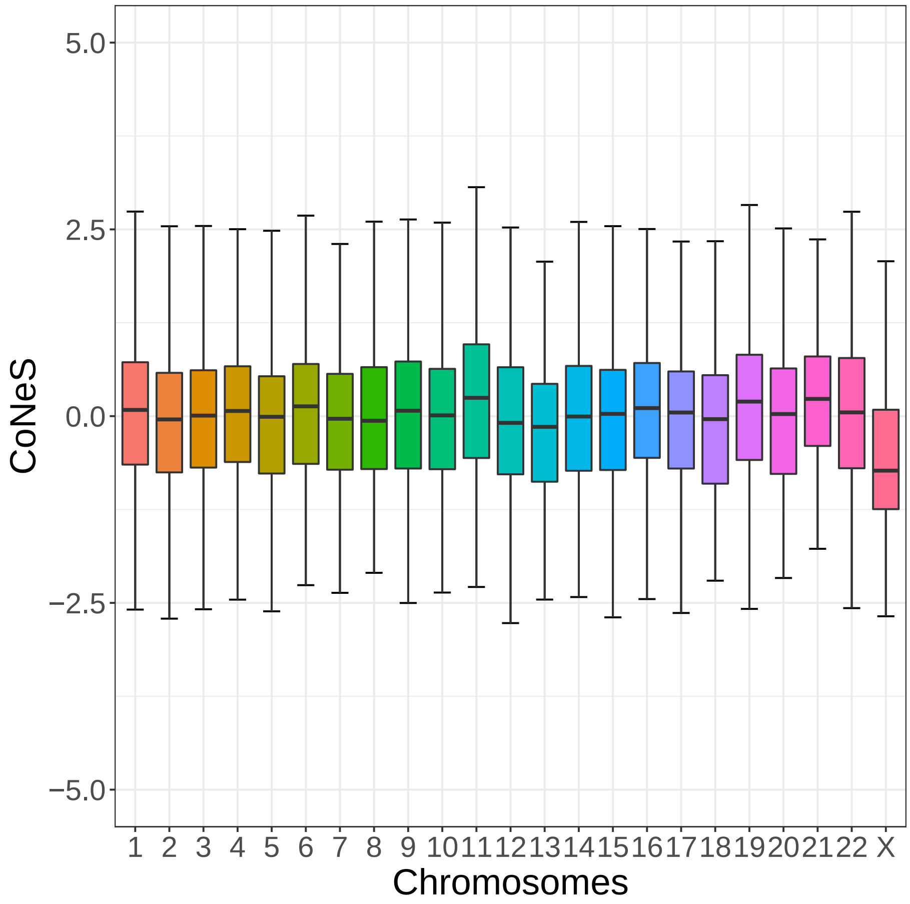


Figure S2: Distribution of CoNeS for each human chromosome.

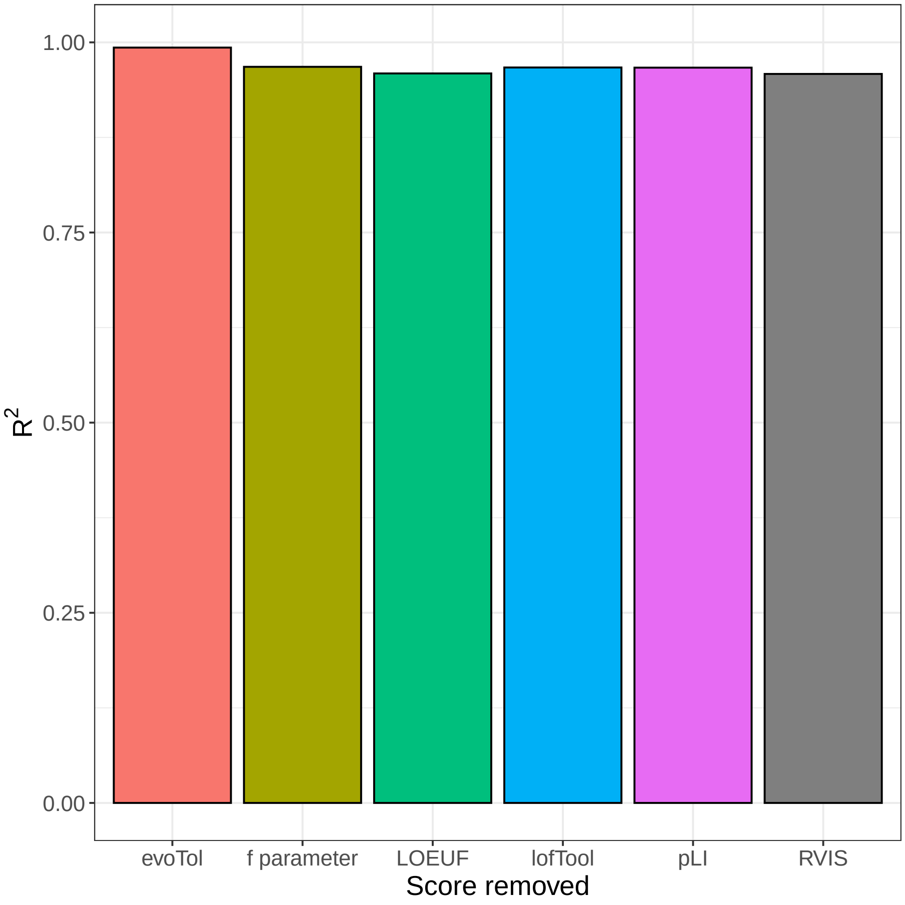


Figure S3: Correlation between CoNeS and a CoNeS score calculated based on all individual scores except one, as measured by Spearman's $R^2$.

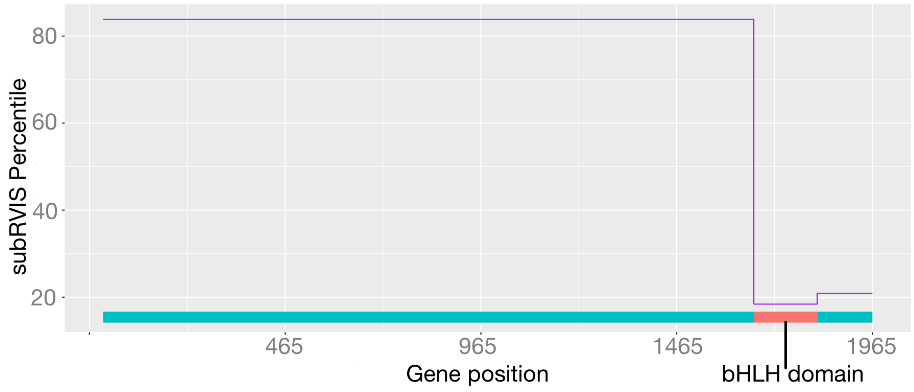


Figure S4: Intolerance to functional variation along the *TCF3* genic region, as measured by subRVIS percentiles.
