## Supplemental Tables for "Negative selection on human genes causing severe inborn errors depends on disease outcome and both the mode and mechanism of inheritance"

| Score | Wilcoxon-based test | Resampling-based test |
|---|---|---|
| evoTol | $3.35 \times 10^{-65}$ | $< 10^{-5}$ |
| $f$ parameter | $5.35 \times 10^{-10}$ | $< 10^{-5}$ |
| LOEUF | $3.36 \times 10^{-13}$ | $< 10^{-5}$ |
| lofTool | $1.64 \times 10^{-66}$ | $< 10^{-5}$ |
| pLI | $2.25 \times 10^{-9}$ | $< 10^{-5}$ |
| RVIS | $1.35 \times 10^{-9}$ | $< 10^{-5}$ |
| SIS | $2.36 \times 10^{-9}$ | $< 10^{-5}$ |
| CoNeS | $2.05 \times 10^{-31}$ | $< 10^{-5}$ |

Table S1: **Statistical significance of differences in negative selection scores between hOMIM genes causing autosomal dominant disease and autosomal background genes.** The $P$-values for a one-tailed Wilcoxon test and a resampling-based test (Methods) assessing the difference between AD hOMIM genes and AB genes are shown.

| Score | Wilcoxon-based test | Resampling-based test |
|---|---|---|
| evoTol | $1.28 \times 10^{-75}$ | $< 10^{-5}$ |
| $f$ parameter | 1 | 0.996 |
| LOEUF | 1 | 1 |
| lofTool | $1.53 \times 10^{-42}$ | $< 10^{-5}$ |
| pLI | 1 | 1 |
| RVIS | 0.484 | 0.661 |
| SIS | 0.990 | 0.990 |
| CoNeS | 0.102 | $1.74 \times 10^{-2}$ |

Table S2: **Statistical significance of differences in negative selection scores between hOMIM genes causing autosomal recessive disease and autosomal background genes.** The $P$-values for a one-tailed Wilcoxon test and a resampling-based test (Methods) assessing the difference between AR hOMIM genes and AB genes are shown.

| Score | Wilcoxon-based test | Resampling-based test |
|---|---|---|
| evoTol | $4.85 \times 10^{-12}$ | $< 10^{-5}$ |
| $f$ parameter | $7.12 \times 10^{-3}$ | $9.62 \times 10^{-2}$ |
| LOEUF | $2.18 \times 10^{-6}$ | $5.97 \times 10^{-3}$ |
| lofTool | $1.75 \times 10^{-12}$ | $< 10^{-5}$ |
| pLI | $3.78 \times 10^{-12}$ | $< 10^{-5}$ |
| RVIS | $8.82 \times 10^{-2}$ | 0.242 |
| CoNeS | $1.08 \times 10^{-12}$ | $< 10^{-5}$ |

Table S3: **Statistical significance of differences in negative selection scores between hOMIM genes causing X-linked disease and X-chromosome background genes.** The $P$-values for a one-tailed Wilcoxon test and a resampling-based test (Methods) assessing the difference between X hOMIM genes and XB genes are shown.

| Score | Wilcoxon-based test | Resampling-based test |
|---|---|---|
| evoTol | $6.98 \times 10^{-3}$ | 0.144 |
| $f$ parameter | 0.806 | 0.933 |
| LOEUF | $3.82 \times 10^{-2}$ | 0.112 |
| lofTool | $1.26 \times 10^{-3}$ | $5.59 \times 10^{-2}$ |
| pLI | 0.932 | 0.967 |
| RVIS | 0.164 | 0.416 |
| SIS | 0.490 | 0.683 |
| CoNeS | $5.95 \times 10^{-2}$ | 0.180 |

Table S4: **Statistical significance of differences in negative selection scores between IEI genes causing autosomal recessive disease and autosomal background genes.** The $P$-values for a one-tailed Wilcoxon test and a resampling-based test (Methods) assessing the difference between AR IEI genes and AB genes are shown.

| Score | Wilcoxon-based test | Resampling-based test |
|---|---|---|
| evoTol | 0.133 | $6.47 \times 10^{-2}$ |
| $f$ parameter | 0.156 | 0.352 |
| LOEUF | $4.14 \times 10^{-2}$ | 0.107 |
| lofTool | $7.09 \times 10^{-3}$ | $3.15 \times 10^{-2}$ |
| pLI | 0.199 | 0.189 |
| RVIS | 0.466 | 0.475 |
| SIS | 0.683 | 0.945 |
| CoNeS | $6.40 \times 10^{-2}$ | 0.175 |

Table S5: **Statistical significance of differences in negative selection scores between IEI genes causing both autosomal dominant and recessive disease and autosomal background genes.** The $P$-values for a one-tailed Wilcoxon test and a resampling-based test (Methods) assessing the difference between AR/AD IEI genes and AB genes are shown.

| Score | Wilcoxon-based score | Resampling-based score |
|---|---|---|
| evoTol | $2.26 \times 10^{-4}$ | $4.5 \times 10^{-3}$ |
| $f$ parameter | $9.41 \times 10^{-2}$ | 0.371 |
| LOEUF | $4.18 \times 10^{-3}$ | 0.183 |
| lofTool | $5.95 \times 10^{-5}$ | $7.60 \times 10^{-2}$ |
| pLI | $1.07 \times 10^{-5}$ | $2.17 \times 10^{-2}$ |
| RVIS | 0.224 | 0.498 |
| CoNeS | $1.19 \times 10^{-5}$ | $2.21 \times 10^{-2}$ |

Table S6: **Statistical significance of differences in negative selection scores between IEI genes causing X-linked recessive disease and X-chromosome background genes.** The $P$-values for a one-tailed Wilcoxon test and a resampling-based test (Methods) assessing the difference between XR IEI genes and XB genes are shown.

| Score | Wilcoxon-based score | Resampling-based score |
|---|---|---|
| evoTol | $1.08 \times 10^{-3}$ | $6.71 \times 10^{-2}$ |
| $f$ parameter | 0.332 | 0.665 |
| LOEUF | $3.10 \times 10^{-3}$ | $9.91 \times 10^{-2}$ |
| lofTool | $8.58 \times 10^{-5}$ | $2.19 \times 10^{-2}$ |
| pLI | 0.439 | 0.667 |
| RVIS | $1.03 \times 10^{-2}$ | $3.25 \times 10^{-2}$ |
| SIS | 0.189 | 0.419 |
| CoNeS | $1.65 \times 10^{-3}$ | 0.112 |

Table S7: **Statistical significance of differences in negative selection scores between IEI genes causing autosomal recessive disease of high severity and autosomal background genes.** The $P$-values for a one-tailed Wilcoxon test and a resampling-based test (Methods) assessing the difference between "high-severity" AR IEI genes and AB genes are shown.

| Score | Wilcoxon-based test |
|---|---|
| evoTol | 0.170 |
| $f$ parameter | 0.130 |
| LOEUF | $1.84 \times 10^{-4}$ |
| lofTool | $3.00 \times 10^{-2}$ |
| pLI | $6.27 \times 10^{-4}$ |
| RVIS | $1.84 \times 10^{-2}$ |
| SIS | $2.25 \times 10^{-2}$ |
| CoNeS | $7.34 \times 10^{-4}$ |

Table S8: **Statistical significance of differences in negative selection scores between genes causing autosomal dominant (AD) IEI by haploinsufficience (HI) and genes causing other AD IEI.** The P-values for a one-tailed Wilcoxon test assessing the difference between HI from gain-of-function (GOF) and dominant negative (DN) genes for each individual score are shown.

| Score | AD | ARAD | Disease severity | CDS length | GC-content |
|---|---|---|---|---|---|
| evoTol | $7.9 \times 10^{-2}$ | 0.21 | 0.35 | $1.8 \times 10^{-6}$ | $2.0 \times 10^{-2}$ |
| $f$ | 0.95 | 0.38 | 0.37 | $-1.5 \times 10^{-5}$ | $1.0 \times 10^{-2}$ |
| LOEUF | 0.61 | 0.22 | 0.30 | $-4.6 \times 10^{-5}$ | $3.9 \times 10^{-4}$ |
| lofTool | 0.57 | 0.40 | 0.34 | $-1.1 \times 10^{-5}$ | $2.6 \times 10^{-2}$ |
| pLI | 1.1 | 0.65 | 0.29 | $-4.4 \times 10^{-5}$ | $2.7 \times 10^{-3}$ |
| RVIS | 0.56 | 0.11 | 0.23 | $1.2 \times 10^{-5}$ | $-2.5 \times 10^{-2}$ |
| SIS | 0.64 | $-2.3 \times 10^{-2}$ | 0.27 | $1.8 \times 10^{-5}$ | $5.6 \times 10^{-3}$ |
| CoNeS | 0.94 | 0.37 | 0.39 | $-1.7 \times 10^{-5}$ | $4.9 \times 10^{-3}$ |

Table S9: **Differences in negative selection scores between IEI-causing genes, according to disease mode of inheritance, clinical severity, gene length and GC content.** This table presents, for each of the individual scores, the effect sizes from a multiple regression model that predicts each negative selection individual score for each autosomal IEI gene. Predictors include AD and ARAD, two binary variables that code if the gene causes AD disease or both AR and AD diseases, respectively, disease severity (high or mild), the coding sequence (CDS) length (in bp) and the CDS content in C and G bases (GC-content).

| Score | Wilcoxon-based score | Resampling-based score |
|---|---|---|
| evoTol | $1.10 \times 10^{-6}$ | $10^{-5}$ |
| $f$ parameter | $6.27 \times 10^{-93}$ | $< 10^{-5}$ |
| LOEUF | $7.12 \times 10^{-97}$ | $< 10^{-5}$ |
| lofTool | $5.93 \times 10^{-74}$ | $< 10^{-5}$ |
| pLI | $4.64 \times 10^{-105}$ | $< 10^{-5}$ |
| RVIS | $8.45 \times 10^{-99}$ | $< 10^{-5}$ |
| SIS | $3.31 \times 10^{-92}$ | $< 10^{-5}$ |
| CoNeS | $4.54 \times 10^{-133}$ | $< 10^{-5}$ |

Table S10: **Statistical significance of differences in negative selection scores between genes causing autosomal dominant inborn errors of neurodevelopment and autosomal background genes.** The $P$-values for a one-tailed Wilcoxon test and a resampling-based test (Methods) assessing the difference between AD IEND genes and AB genes are shown.

| Score | Wilcoxon-based score | Resampling-based score |
|---|---|---|
| evoTol | $1.30 \times 10^{-7}$ | $10^{-5}$ |
| $f$ parameter | $2.32 \times 10^{-2}$ | 0.199 |
| LOEUF | 0.920 | 0.996 |
| lofTool | $2.33 \times 10^{-4}$ | $9.16 \times 10^{-2}$ |
| pLI | 1 | 1 |
| RVIS | $3.66 \times 10^{-14}$ | $< 10^{-5}$ |
| SIS | $8.86 \times 10^{-6}$ | $5.41 \times 10^{-2}$ |
| CoNeS | $3.69 \times 10^{-4}$ | 0.193 |

Table S11: **Statistical significance of differences in negative selection scores between genes causing autosomal recessive inborn errors of neurodevelopment and autosomal background genes.** The $P$-values for a one-tailed Wilcoxon test and a resampling-based test (Methods) assessing the difference between AR IEND genes and AB genes are shown.

| Score | Wilcoxon-based score | Resampling-based score |
|---|---|---|
| evoTol | $9.11 \times 10^{-6}$ | $< 10^{-5}$ |
| $f$ parameter | $3.49 \times 10^{-9}$ | $< 10^{-5}$ |
| LOEUF | $4.49 \times 10^{-8}$ | $1.7 \times 10^{-4}$ |
| lofTool | $2.48 \times 10^{-11}$ | $< 10^{-5}$ |
| pLI | $4.50 \times 10^{-4}$ | $2.8 \times 10^{-3}$ |
| RVIS | $8.03 \times 10^{-12}$ | $< 10^{-5}$ |
| SIS | $8.98 \times 10^{-8}$ | $5 \times 10^{-5}$ |
| CoNeS | $4.55 \times 10^{-14}$ | $< 10^{-5}$ |

Table S12: **Statistical significance of differences in negative selection scores between genes causing both autosomal dominant and recessive inborn errors of neurodevelopment and autosomal background genes.** The $P$-values for a one-tailed Wilcoxon test and a resampling-based test (Methods) assessing the difference between AR/AD IEND genes and AB genes are shown.

| Score | Wilcoxon-based score | Resampling-based score |
|---|---|---|
| evoTol | $4.71 \times 10^{-2}$ | $2.17 \times 10^{-2}$ |
| $f$ parameter | $1.07 \times 10^{-20}$ | $< 10^{-5}$ |
| LOEUF | $8.77 \times 10^{-17}$ | $< 10^{-5}$ |
| lofTool | $1.58 \times 10^{-14}$ | $< 10^{-5}$ |
| pLI | $4.48 \times 10^{-31}$ | $< 10^{-5}$ |
| RVIS | $5.44 \times 10^{-18}$ | $< 10^{-5}$ |
| CoNeS | $3.40 \times 10^{-37}$ | $< 10^{-5}$ |

Table S13: **Statistical significance of differences in negative selection scores between genes causing X-linked inborn errors of neurodevelopment and X-chromosome background genes.** The $P$-values for a one-tailed Wilcoxon test and a resampling-based test (Methods) assessing the difference between X IEND genes and XB genes are shown.

| Score | Separation between HI and not HI IEND genes |
|---|---|
| evoTol | 0.737 |
| $f$ parameter | $2.29 \times 10^{-2}$ |
| LOEUF | $1.73 \times 10^{-13}$ |
| lofTool | $3.19 \times 10^{-6}$ |
| pLI | $2.63 \times 10^{-19}$ |
| RVIS | $9.05 \times 10^{-8}$ |
| SIS | $4.58 \times 10^{-5}$ |
| CoNeS | $1.12 \times 10^{-13}$ |

Table S14: **Statistical significance of differences in negative selection scores between genes causing autosomal dominant (AD) inborn errors of neurodevelopment (IEND) by haplo-insufficiency and other AD IEND genes.** The $P$-values for a one-tailed Wilcoxon test assessing the difference between HI AD IEND genes and non-HI AD IEND genes are shown.
